# Decrease in Microbial Diversity Along a Pollution Gradient in Citarum River Sediment

**DOI:** 10.1101/357111

**Authors:** Anniek de Jong, Bas van der Zaan, Gertjan Geerlings, Michiel in’t Zandt, Lufiandi, Dwina Roosmini, Herto Dwi Ariesyady, Claudia Lüke, Mike Jetten

## Abstract

Pollution of water resources is a major risk to human health and water quality throughout the world. Here, we studied the effect of river pollution on the microbial community composition in the Citarum River Basin, West Java, Indonesia. Sediment was collected at six sampling points along a gradient of pollution, from the pristine source of the river to the heavily polluted downstream site in the densely populated urban area of Bandung. After DNA extraction, microbial diversity and potential for nitrogen cycling were analyzed based on 16S rRNA gene amplicon sequencing and metagenomics. Comparing the pristine sediment to the polluted site showed a lower microbial diversity and higher dominance of anaerobic processes than at the polluted site. The most dominant phylum within the Bacteria were the *Proteobacteria*, which shifted from *Beta*- and *Deltaproteobacteria* in the pristine site to *Alpha*- and *Gammaproteobacteria* in the polluted sediment. With pollution *Actinobacteria, Bacteroidetes,* and *Firmicutes* increased in relative abundance. Relative high abundance of *Soil Crenarchaeotic group* was found in all sample sites, whilst the methanogenic archaea became more abundant with increased pollution and anaerobicity. The study of the nitrogen cycling potential revealed that ammonium oxidation and denitrification appeared to be abundant processes in the pristine site, whereas ammonification seemed to be more important in the polluted site. Increased water treatment would restore water quality and microbial diversity in Citarum river sediments.

## INTRODUCTION

Rapid economic development, industrialization, urbanization and population growth have seriously impacted river quality and contributed to river pollution. Contamination of water resources is a major risk to water quality and ecological functioning of rivers, and poses serious threats to the health of the human population and livestock inhabiting the catchment area. Pollution is one of the most obvious environmental threats to the ecological health of rivers and streams and could strongly influence the diversity of the microorganisms living in the sediment. The microbial abundance and functional diversity is strongly affected by environmental factors (Allison and Martiny, 2008; Sun *et al.*, 2012). The variance in bacterial community correlated most strongly with water temperature, conductivity, pH and dissolved oxygen (DO) in freshwater, intertidal wetland and marine sediment (Wang *et al.*, 2012). Additionally, microbial diversity was decreased by environmental pollution in a tropical river (Kochling *et al.*, 2017). Increase in river metal content was correlated with higher abundance of Alphaproteobacteria in freshwater sediment (Zhu *et al.*, 2013). Furthermore, chemicals from textile and dying industry process waters precipitate in river sediment where they are degraded by microbial community on the order of years, thus inducing large changes in the microbial diversity (Ito *et al.*, 2016).

Here, we focus on the Citarum river located in West Java, Indonesia, which originates at Mount Wayang from where it runs to the northern coast of Java close to Jakarta. The Citarum river has an important role in life of the people in the catchment area, with a population of 40 million. It is a source of water for agriculture, industry, domestic water supply and for disposal of sewage. The river is also important for generating of electricity by three hydropower plants, delivers 20% of Indonesia’s gross domestic product and supplies drinking water to 80% of the population of Jakarta (10 million people) (United Nations Department of Economic and Social Affairs, 2017). Given these important functions, Citarum is categorized as a ‘super priority river’ by the government in 1984, nevertheless the Citarum river is considered one of the most polluted rivers in the world. When Indonesia experienced a manufacturing boom in the textile industry, waste water disposal was not a high priority. As a result, the Citarum showed severe visible problems of garbage disposal, and untreated industrial and domestic wastewater discharge (Fulazzaky *et al.*, 2008). Similarly, the waste of agriculture is disposed untreated into the river, and thus tons of cattle manure ends up in the water each day. Together these pollutions have resulted in more than 60% reduction of the fish, so the catch-of-the-day is nowadays old plastic that is sold for recycling. In the upstream area wide spread deforestation to accommodate fields for vegetables have led to erosion and land degradation (Ministry of Public Works, 2007). High sedimentation generates a decline of the drainage area causing floods, whereby more people come in contact with the highly polluted river water. Current problems faced by Citarum river are very complex and a simple solution is insufficient to solve the multitude of problems. In 2018, President of Indonesia formulated the ambition to make the water in Indonesia’s most strategic river basin drinkable by 2025.

The aim of present study was to gain a better understanding of the sediment associated microbial diversity change and response to the environmental pollution in the Citarum river. Sediment samples were collected along a gradient of pollution, from the source of the river to the heavily polluted downstream urban area, distributed over six sampling points. Microbial diversity and nitrogen gene functionality were analyzed based on 16S rRNA gene amplicon sequencing and metagenomics.

## MATERIALS AND METHODS

### Sampling sites

The Citarum river is the largest river in West Java Province, Indonesia, with a catchment area of approximately 6867 km^2^ and a length of 269 km (Sellmann and Bogner, 2012a). The river originates in the area surrounding Mount Wayang from where it runs to the northern coast of Java. It flows the first 200 km through mountainous and hilly terrain, followed by three cascade reservoirs, from where it travels through a plain area for 70 km where it drains into the Java sea, east of Jakarta (Figure 1). The climate is characterized by two distinct seasons: rainy season (November to April) and dry season (May to October) (Ministry of Public Works, 2007). The annual rainfall in the Citarum watershed is about 2580 mm, with approximately 1840 mm in the rainy season and around 740 mm in the dry season. The stream flow is highly variable between the seasons, but the average flow is 173 m^3^/s (Sellmann and Bogner, 2012b). Flooding is a common occurrence, especially during the rainy season.

Sediment samples were taken in June 2014 at six different (named S1 – S6) locations in the Citarum river (Figure 1, Table 1), from the source of the Citarum river where the water is pristine till downstream the city of Bandung where the water is highly polluted. At each site, five sediment samples were taken, each of them composed from different sub-samples. From S1, S2, S4 and S6 sediment was taken from the shoreline and the middle of the river, whereas from S3 and S5 samples could only be taken from the shoreline. Sediment samples were collected in sterile 50 mL centrifuge tubes (VWR, Amsterdam, The Netherlands). Field parameters consisting of electrical conductivity, pH, temperature, total dissolved solids and dissolved oxygen were taken at each sample location (Table 2). Samples were transported on ice, where after samples were stored in the freezer.

**FIGURE 1.**
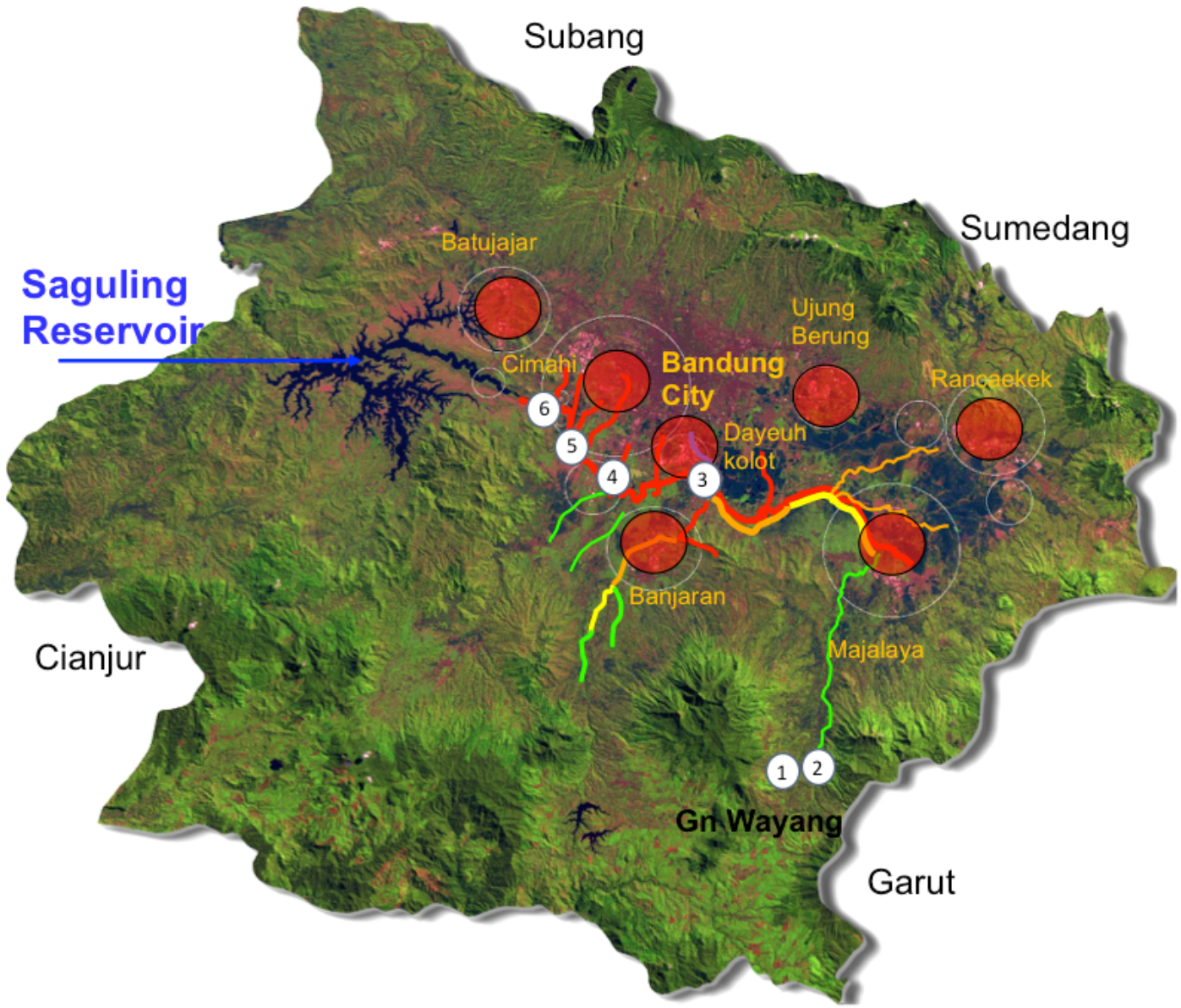
Map of the Upper Citarum River sampling sites. With circles - sampling sites from current study; Red circles - industrial zone; Quality of river water in colors: green - good (DO>3 ppm), yellow - bad (2<DO<3 ppm), red - extremely polluted (DO<2 ppm).

**Table 1.**
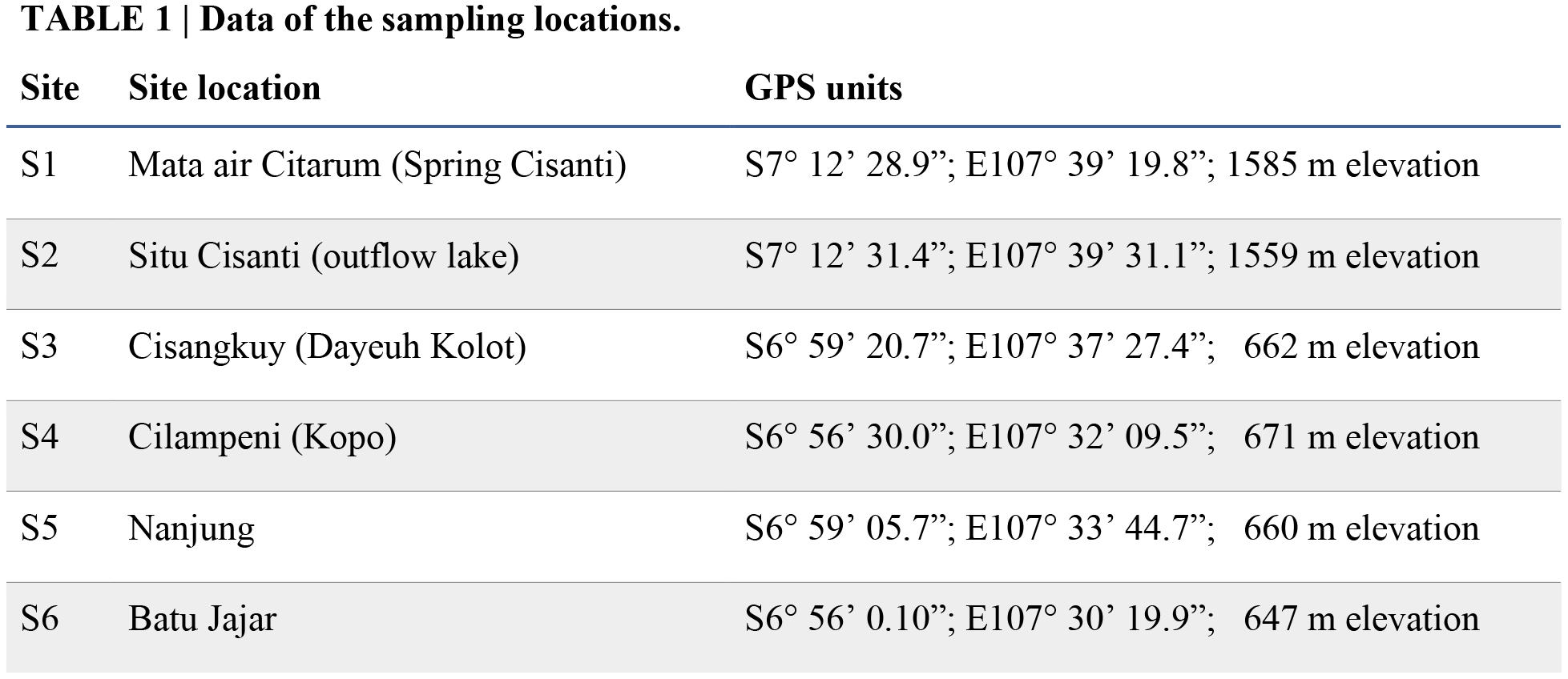
Data of the sampling locations.

### DNA extraction

DNA extraction from each of the five samples at each site was performed using two different methods. DNA extraction using the Powersoil DNA Isolation Kit (MO BIO, Qiagen, Venlo, the Netherlands) was performed according the manufacturer’s instructions with the following modifications: for homogenization of the sample a bead beater was used at 30 bps for 2 minutes. DNA was eluted in 50 μl diethylpyrocarbonate-treated nuclease-free water (DEPC) after incubation of 5 minutes. The second DNA isolation using CTAB/Phenol/Chloroform was done as described in Speth *et al.* (2016). Followed by DNA purification over a column provided in the Powersoil^®^ DNA isolation kit (follow manufacture’s protocol from the addition of Solution C4). DNA quality was checked by agarose gel electrophoresis, spectrophotometrically using the NanoDrop 1000 (Invitrogen, Thermo Fisher, Carlsbad CA, USA) and fluorometrically using the Qubit dsDNA HS Assay Kit (Invitrogen, Thermo Fisher, Carlsbad CA, USA) according to manufacturer’s instructions. DNA extracted by the different methods was pooled per sediment sample.

### 16S rRNA gene amplicon sequencing and analysis

Bacteria and Archaea16S rRNA gene amplicon sequencing was performed on an Illumina Miseq Next Generation Sequencing by Macrogen, Korea. The primers for bacterial amplification were Bac341F 5‘- CCTACGGGNGGCWGCAG-3’) (Herlemann *et al.*, 2011) and Bac806R (5‘-GGACTACHVGGGTWTCTAAT-3’) (Caporaso *et al.*, 2012). For archaeal amplification the primers were Arch349F (5‘-GYGCASCAGKCGMGAAW-3’) and Arch806R (5‘-GGACTACVSGGGTATCTAAT-3’)(Takai and Horikoshi, 2000). The output data had a Q20 quality score … % for …% of the reads or higher (Q30 for …% of the reads). Average total read count was…. reads per sample. Length of the sequences used for analyses was between 400 and 500 base pairs (bp). For additional single-end read analysis minimum read length was set to 291 bp. The UCHIME algorithm (Edgar *et al.*, 2011) was used to checked for and discard chimeras. Thereafter reads were clustered by OTU with a 97% identity cut-off value and classified using the SILVA v128 16S rRNA gene non-redundant database (SSURef_NR99_128_SILVA) and the Bayesian classifier (‘wang’) using the MySeq SOP (http://www.mothur.org/) (Schloss *et al.*, 2009). In all datasets ‘Chloroplasts’, ‘Mitochondria’, ‘unknown’ and ‘Eukaryota’ were removed, whereas ‘Bacteria’ and ‘Archaea’ were removed for the archaeal and bacterial (Luesken *et al.*, 2012)(Kirschke *et al.*, 2013; Pjevac *et al.*, 2017) sequence datasets, respectively. For the downstream analysis with R (https://www.r-project.org/) (R Development Core Team, 2013) and Rstudio v3 (https://www.rstudio.com/) (RStudio Team, 2015) the ‘filename.taxonomy’ and ‘filename.shared’ files were used as described in in ‘t Zandt et al. (2017). Singletons were removed and data was rarefied in R (see rarefraction curves in Figure S1). All sequencing data were submitted to the GenBank databases under the BioProject PRJNA478143.

### Metagenome sequencing

All kits described in this paragraph were obtained from Life technologies (Life technologies, Carlsbad, CA, USA). Metagenome sequencing was performed with Ion Torrent Personal Genome Machine (PGM) System for the pristine site (S1) and the highly polluted site (S6). The 10 DNA samples per site were pooled according to concentration. DNA of both samples was sheared for 4 minutes using IonXpress Plus Fragment Library Kit following manufacturer’s instructions and barcoded using the IonXpress barcode adapters. For size selection, the E-Gel SizeSelect 2% agarose gel was used. Emulsion PCR was performed using OneTouch 400bp kit and sequencing was done on an IonTorrent PGM using the Ion PGM 400bp sequencing kit and an Ion 318v2 chip. The raw reads were submitted to BioProject PRJNA478143. Raw sequence reads were imported int the CLC Genomics Workbench (v7.0.4, CLCbio Arhus, Denmark) and end-trimmed on quality using the CLC genomics default settings (quality limit 0.05 and two ambiguous nucleotides allowed) and length (≥100 bp) resulting in X million reads for S1 and X million reads from S6. 16S rRNA gene analysis and functional gene (amoA, narG, napA, nas, cbbL, mcrA and pmoA) analysis were done as described in Lüke et al. (2016). Read counts of the selected marker genes were normalized to gene length and total read abundance in the metagenome dataset.

## RESULTS

### Site description and sediment sampling

Physical and chemical characteristics of the Citarum river samples such as temperature and pH, conductivity, dissolved oxygen (DO) and total dissolved solids (TDS) were determined at every sample location (Table 2). The temperature values for water samples ranged from 20°C in Mata Air Citarum (S1) to 27°C at Cilampeni (S4). The pH was at all sites between 6 and 7. Furthermore, the conductivity of the water increased as the river got more contaminated. The DO was different at each site, Situ Cisanti (S1) had the highest amount of oxygen in the water followed by Cilampeni (S4). At both sites the water flow was high, which resulted in more oxygen in the water. The TDS increased with the pollution but did not exceed the maximum contamination level of 500 ppm advised by the EPA Secondary Regulations.

**Table 2.**
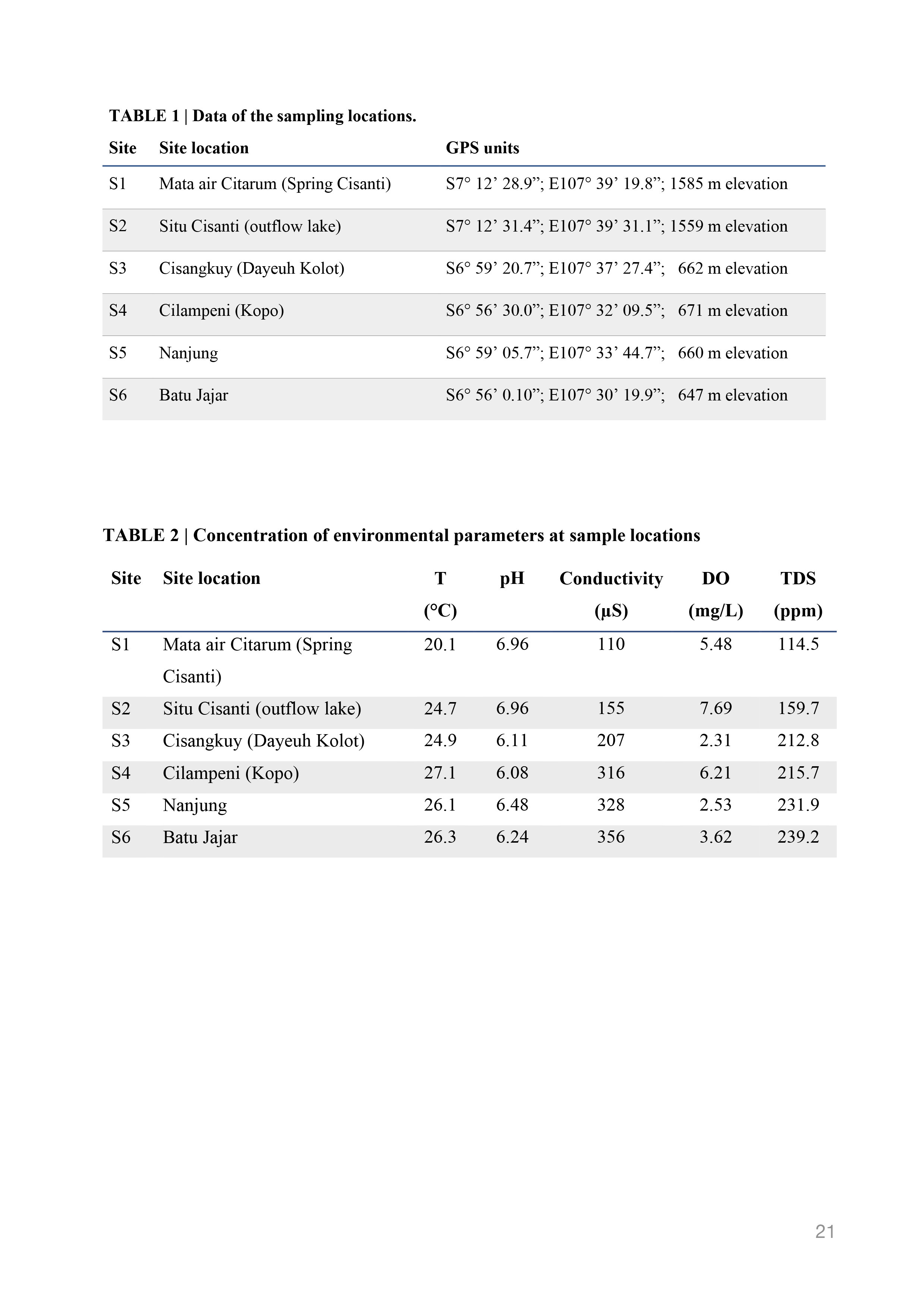
Archaeal alpha diversity analyses. All data sets were rarefied for the smallest dataset = 3,905 sequences. Sample coverages were all above 91 %.

### Microbial community structure based on 16S rRNA gene amplicon sequencing

For each site, the 16S rRNA genes were amplified using two primers sets targeting Bacteria and Archaea, respectively. The Shannon and Simpson diversity analyses showed a high bacterial and archaeal diversity at the source of the river (S1 and S2), however with increasing pollution the microbial diversity decreased (Table 3 and 4). The Shannon evenness showed a decrease in evenness along the gradient of pollution, with highest evenness found in the pristine site, however, this was not visible from the Simpson indexes (Table 3 and 4). No trend could be found in the Chao1 species richness estimations (Table 3 and 4), indicating that the number of species is similar per site, which was also reflected by the non-metric multidimensional scaling (NMDS) plots (SFigure 1 and 2).

**Table 3.** Geometric overlap between EB tiles and wedges.

| Sample | Sampling coverage | Diversity |  | Richness | Evenness |  |
| --- | --- | --- | --- | --- | --- | --- |
|  |  | Simpson index | Shannon index | Chao1 estimator | Simpson-based evenness | Shannon-based evenness |
| Cit_S1 | 0.91 | 0.03 | 4.81 | 1275 | 0.06 | 0.75 |
| Cit_S2 | 0.91 | 0.03 | 4.57 | 1305 | 0.07 | 0.73 |
| Cit_S3 | 0.93 | 0.04 | 4.48 | 938 | 0.05 | 0.72 |
| Cit_S4 | 0.93 | 0.05 | 4.28 | 1131 | 0.04 | 0.69 |
| Cit_S5 | 0.93 | 0.04 | 4.38 | 942 | 0.05 | 0.70 |
| Cit_S6 | 0.94 | 0.04 | 4.22 | 1029 | 0.06 | 0.69 |
| Average | 0.93 | 0.04 | 4.46 | 1103 | 0.06 | 0.71 |
| Stdev | 0.01 | 0.01 | 0.22 | 161 | 0.01 | 0.02 |

**Table 4.** Bacterial alpha diversity analyses. All data sets were rarefied for the smallest dataset = 36,666 sequences. Sample coverages were all above 63%.

| Sample | Sampling coverage | Diversity |  | Richness | Evenness |  |
| --- | --- | --- | --- | --- | --- | --- |
|  |  | Simpson index | Shannon index | Chao1 estimator | Simpson-based evenness | Shannon-based evenness |
| Cit_S1 | 0.63 | 0.00 | 8.75 | 72599 | 0.04 | 0.90 |
| Cit_S2 | 0.70 | 0.01 | 7.98 | 63320 | 0.01 | 0.84 |
| Cit_S3 | 0.69 | 0.01 | 7.95 | 67977 | 0.01 | 0.83 |
| Cit_S4 | 0.67 | 0.00 | 8.03 | 79644 | 0.02 | 0.84 |
| Cit_S5 | 0.68 | 0.01 | 7.99 | 65837 | 0.01 | 0.83 |
| Cit_S6 | 0.69 | 0.01 | 7.78 | 72462 | 0.01 | 0.82 |
| Average | 0.68 | 0.01 | 8.08 | 70306 | 0.01 | 0.84 |
| Stdev | 0.02 | 0.00 | 0.34 | 5853 | 0.01 | 0.03 |

Forty to 60% of the bacterial community structure was made up of reads affiliated to groups with an individual abundance of less than 2%. The rare microbial biosphere decreased in S3-S6, but was still around 25%. *Proteobacteria* were the most dominant phylum in the river sediment (Figure 2), with the relative abundance between 21-38% at the different sites. Beta- and *Deltaproteobacteria* were the most abundant classes in the sediment of the source of river (S1 and S2) (with 15-18% *Betaproteobacteria* 8-11% *Deltaproteobacteria),* followed by *Alphaproteobacteria* (3-9%) and *Gammaproteobacteria* (3%). The polluted sediment was dominated by *Alphaproteobacteria* (between 11-13%), followed by Beta- and Gamma- classes, and only 4% of reads affiliated to *Deltaproteobacteria* at S4. Deltaproteobacterial reads in S1 and 2 were mostly affiliated to *Desulfuromonadales, Myxococcales* and *Syntrophobacterales*. *Bacteroidetes* were only found in relative high abundance in the polluted sites (8-21%). Most abundant orders were *Flavobacteriales* (2-14%), *Bacteroidales* (2-9%), and *Sphingobacteriales* (2-3%). Interestingly, the *Flavobacteriales* were most abundant at S3, however as soon as the pollution started, they decreased in read numbers till less than 2% of the relative abundance at S6. At S6, the most polluted site, *Lactobacillales* was present with a relative abundance of 15 %. Furthermore, in the sediment of the polluted site *Clostridia* increased in relative abundance compared to the origin of the river (4-8% in S1 and 2 to 14% in S6).

**Figure 2.**
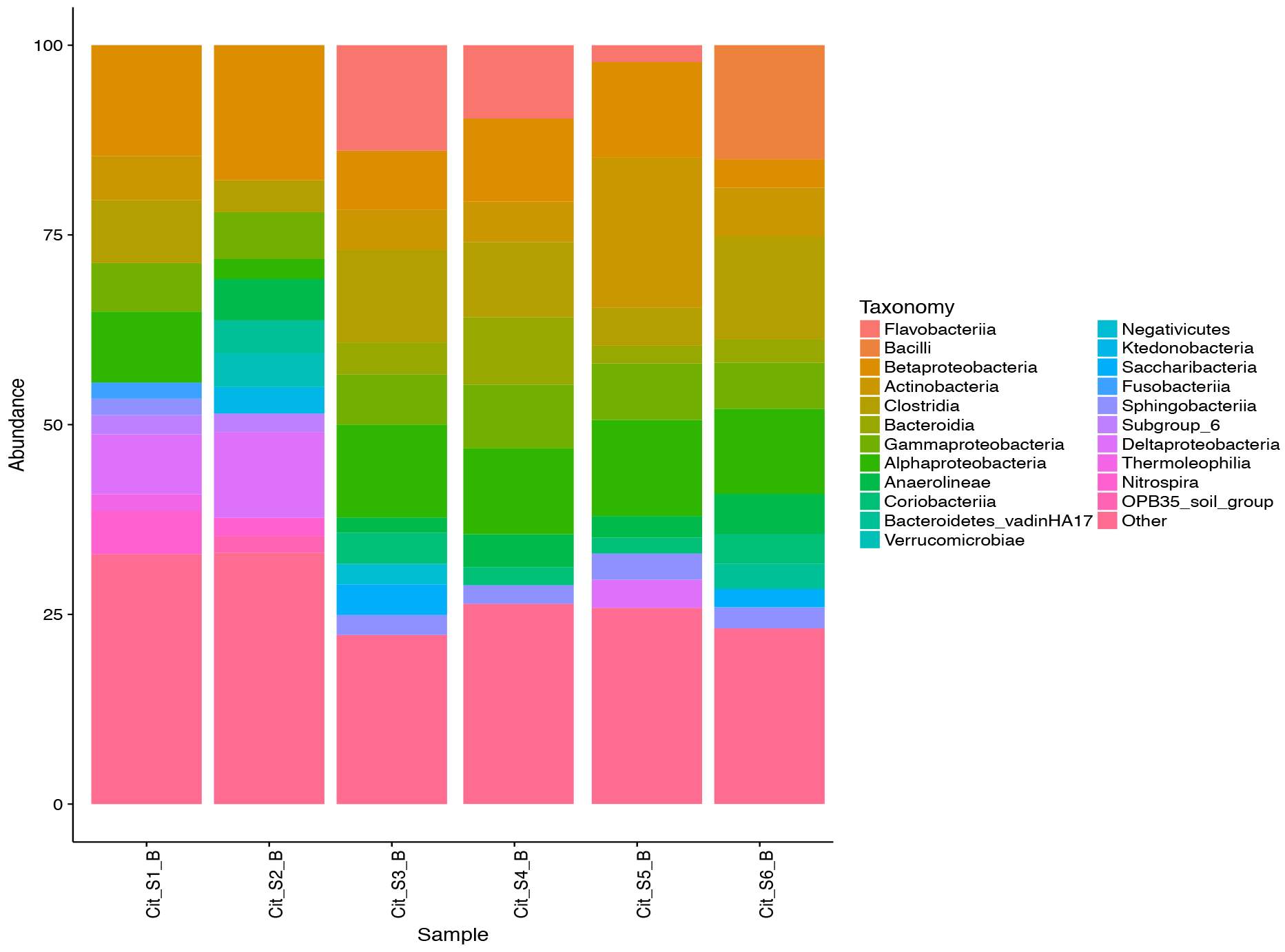
Phylogenetic classification of amplified bacterial 16S rRNA genes in the river sediment from S1 to S6. The maximum taxonomy depth is on class level, whereby taxonomic groups with less than 2% abundance are grouped in others.

The archaeal community structure was also influenced by pollution and became less diverse at the more polluted sites. *Soil Crenarchaeotic group* had the highest relative abundance at all sites (between 28-79%), followed by *Methanosaetaceae* (6-24%) (Figure 3). The methanogens *Methanosarcinaceae* and *Methanobacteriaceae* became more abundant with pollution, while *Bathyarchaeota* (12-15% in S1 and S2) and *Methanoregulaceae* (17% in S2) were mostly present at the origin of the river.

**Figure 3.**
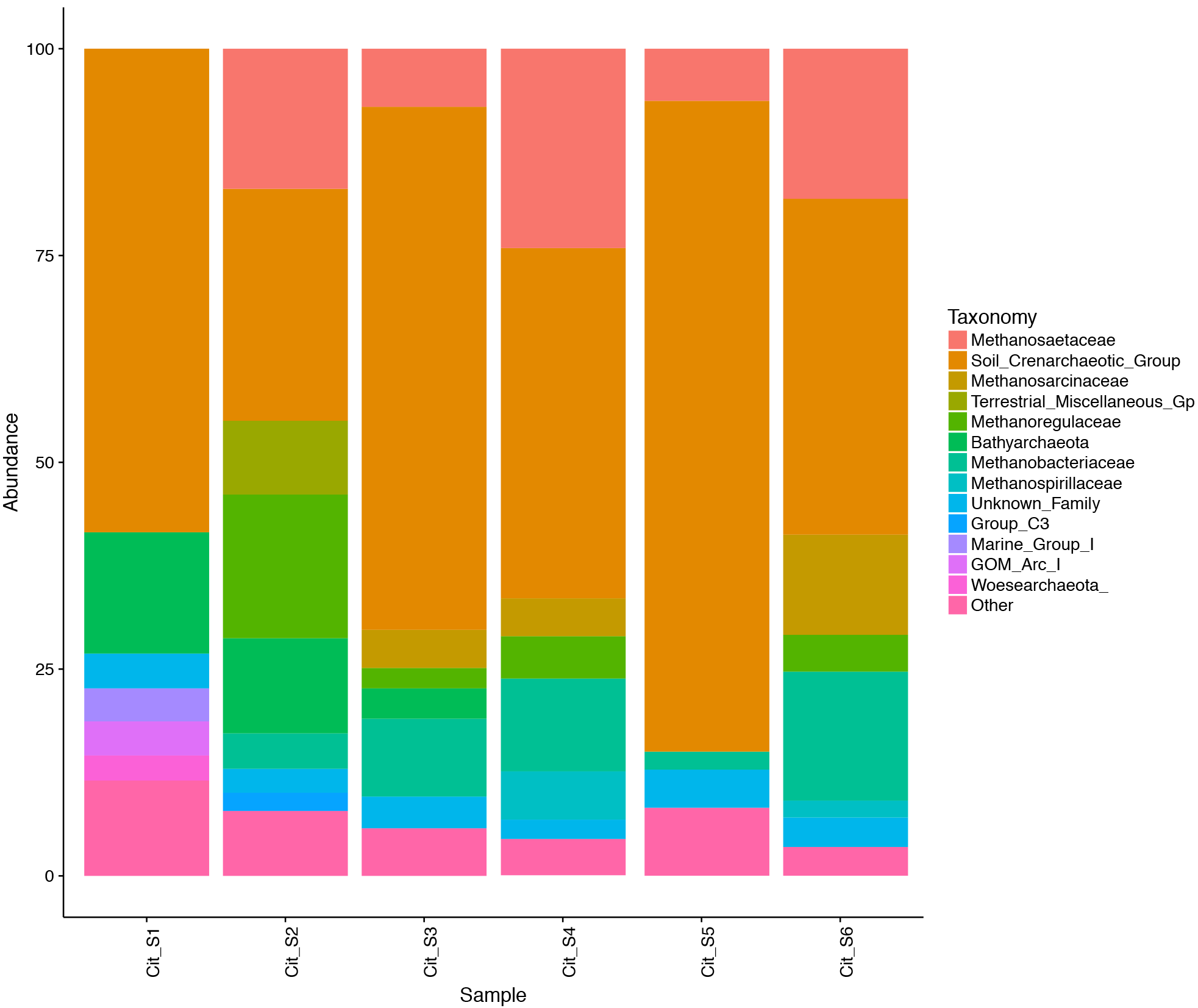
Phylogenetic classification of amplified archaeal 16S rRNA genes in the river sediment from S1 to S6. The maximum taxonomy depth is on family level, whereby taxonomic groups with less than 2% abundance are grouped in others.

### 16S rRNA gene analysis of S1 and S6 by metagenomics

Metagenomic datasets of Citarum river sediment were analyzed for 16S rRNA gene diversity, and functional of nitrogen and methane cycle potential. In total, 5.1 million reads were sequenced (Table S1) with Ion Torrent Technologies. Based on the 16S rRNA gene, a phylogenetic classification could be made. In both sites the bacteria dominated and the abundance of archaea was 3.2% at S1 and 2.7% at S6). The archaea in river sediment had equal amounts of reads affiliated to *Thaumarchaeota* and *Euryarchaeota* at S1, whereas at S6 a relative high abundance of *Euryarchaeota* was observed (Figure 4). Similar to the 16S rRNA amplicon data, the bacterial domain was dominated by *Proteobacteria* in both samples (S1 36% and S6 25%) (Figure 4). The *Beta-* and *Deltaproteobacteria* were more abundant at S1 (13% and 10% at S1 compared to 6% and 2% at S6), while at S6 more reads were affiliated to *Alpha*- (7% versus 8% at S1 and S6) and *Gammaproteobacteria* (7% at S1 and 10% at S6) (Figure 5). The *Betaproteobacteria* at S1 were dominated by the orders *Burkhorderiales* and *Nitrosomonadales*, for S6 this class was dominated by *Burkhorderiales*. *Deltaproteobacteria* were dominated by the orders *Desulfuromonadales* and *Myxococcales* at S1 and contained mostly *Myxococcales* at S6. The *Alphaproteobacteria* in both were samples dominated by the order *Rhizobiales,* followed by *Rhodospirillales* and *Rickettsiales* for S1, and *Rhodobacterales* and *Shingomonadales* for S6. The *Gammaproteobacteria* were more abundant and diverse at S6 with dominating orders *Enterobacteriales* and *Pseudomonadales*. The relative abundance of *Acidobacteria, Cyanobacteria,* and *Nitrospirae* was higher at S1 compared to S6 (6%, 8% and 5% at S1 to 3%, 1% and 0.3% at S6 respectively). In the sediment of the highly polluted site *Actinobacteria* (S1 6% and S6 13%), *Bacteroidetes* (S1 5% and S6 8%), and the *Firmicutes* (S1 5%, S6 20%) had increased in relative abundance.

**Figure 4.**
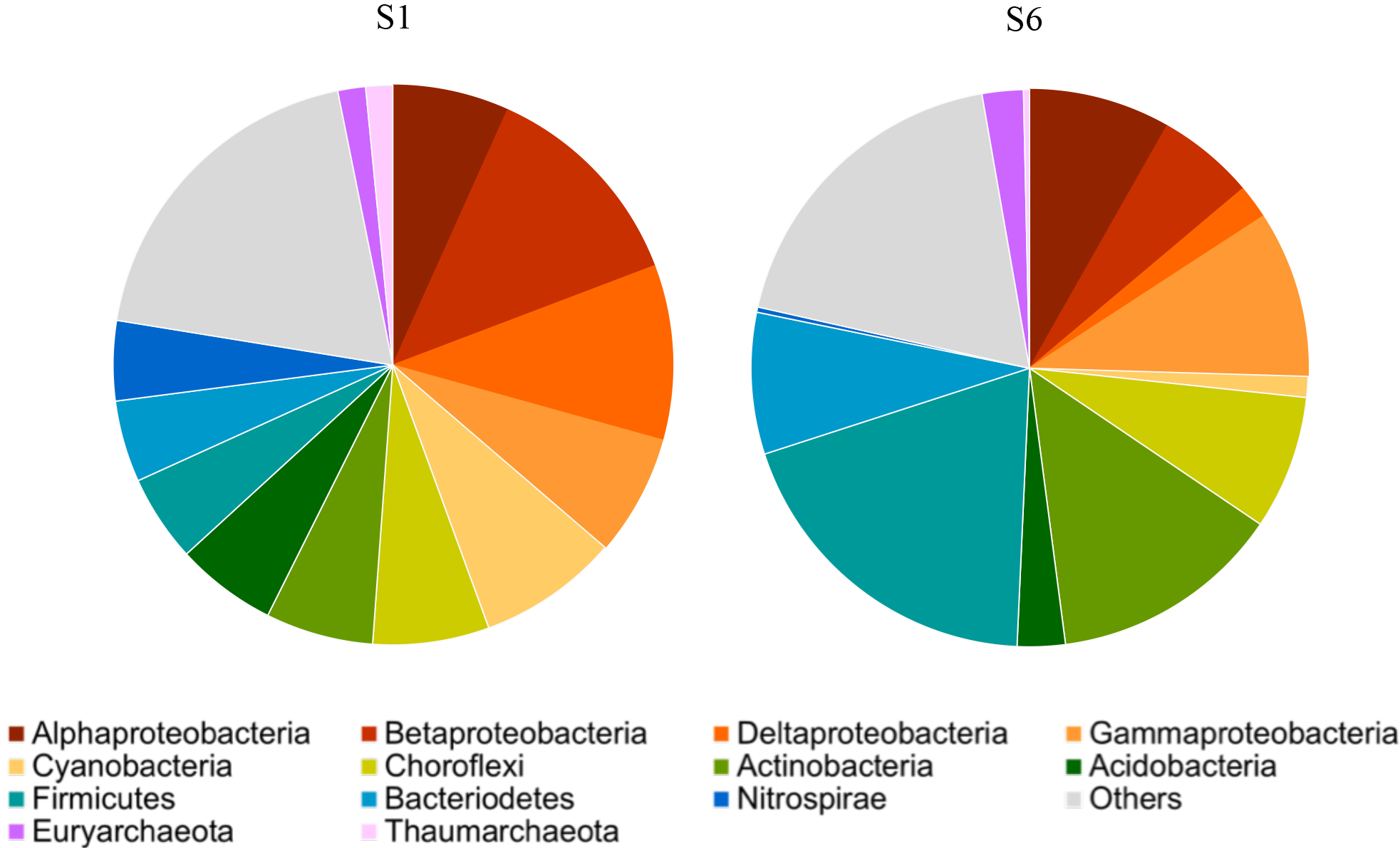
Relative abundances of major phyla of Bacteria and Archaea found in metagenomes of S1 and S6 sediment expressed as percentage of total 16S rRNA gene sequences that could be retrieved.

**Figure 5.**
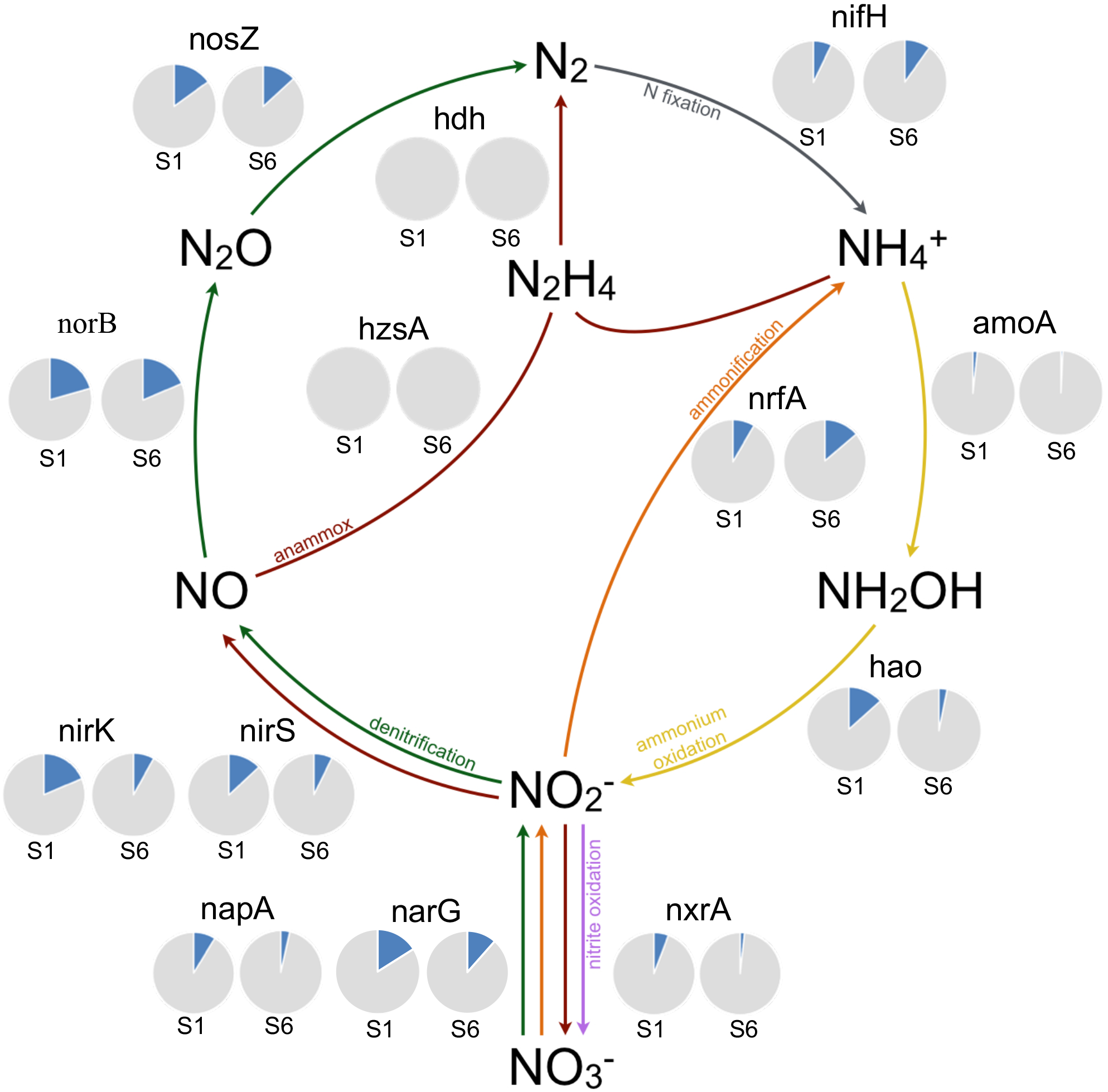
Overview of nitrogen cycle processes by selected marker genes. Read counts were normalized to gene length and total read abundance in the metagenome datasets of S1 and S6 sediments. Normalized read counts are shown as proportion (blue) of total 16S rRNA gene (grey). nifH: nitrogenase, amoA: ammonium monooxygenase, hao: hydroxylamine oxidoreductase, nxrA: nitrate/nitrite oxidoreductase, narG/napA: nitrate reductase, nirS: heme *cd1* copper nitrite reductase, nirK: copper nitrite reductase, norB: nitric oxide reductase, nosZ: nitrous oxide reductase, nrfA: cytochrome c nitrite reductase, hzsA: hydrazine synthese, hdh: hydrazine dehydrogenase.

### Genetic potential for nitrogen and methane cycling

#### Nitrogen cycling potential

To study the nitrogen and methane cycling the nitrogen and methane related functional genes were analyzed to gain more insight how river pollution might influence genomic potential for these processes. We performed Blastx searches against curated key gene databases for nitrogen and methane cycling provided by Lüke *et al.* (2016) (STable X). In general, both sediment samples harbored a wide array of N-cycle genes, covering about a quarter of the abundance of 16S rRNA genes.

The first step in nitrification, from ammonia to nitrite, is encoded by the marker genes *amoA* (ammonium monooxygenase) and *hao* (hydroxylamine monooxygenase). In total 12 reads matched *amoA* in S1, 8 of which could be assigned to ammonia oxidizing archaea (AOA) while the others were assigned to ammonia oxidizing bacteria (AOB). This indicates that the AOA contribute approximately 1.7% of the total microbial community. At the polluted site the assigned *amoA* reads were even lower, 3 reads for AOA and 1 for AOB. The marker gene *hao* was affiliated to 175 reads in S1 and decreased to 35 reads in the polluted site. For S1 94% of the reads were affiliated to Bacteria, 32% of which were assigned to *Proteobacteria* and 16% to *Planctomycetes*. At S6 14 *hao* matches were affiliated with *Proteobacteria* and 5 with *Planctomycetes*.

The second step in nitrification is the oxidation of nitrite to nitrate, which was studied using the marker genes nitrite oxidoreductase (*nxrA*) and nitrate reductase (*narG/napA*). Since *nxrA* and *narG* are homologous enzymes, same method was used as Luke et al (2016) to distinguish between them. In total, 29% and 10% of reads matching *narG/nxrA* were assigned to *nxrA* of *Nitrospira/Nitrospina/anammox* in S1 and S6 respectively, and 2% and 1% of the reads were affiliated to *nxrA* of *Nitrobacter/Nitrococcus/Nitrolancetus* group in S1 and S6, respectively. To further separate the *Nitrospira/Nitrospina/annamox nxrA* classes from S1 MEGAN was used, which showed that 39% of the reads matched to *anammox* and 36% of the reads matched to *Nitrospira*. *Anammox* was not detected with 16S rRNA gene amplicon sequencing and also not found in 16S rRNA gene in the metagenome data. However, after closer inspection, the sequence identity of the assigned *anammox* were all below 80%.

From nitrite there are three pathways possible, denitrification, ammonification and anammox. The distribution of the gene *narG* was diverse and was mostly affiliated to *Proteobacteria* in both sites. The marker gene *napA* was also used to identify nitrate reduction potential in the sediment, which was as *narG*, more found at S1 compared to S6 (163 reads to 56 reads), mostly affiliated to *Proteobacteria.* Nitrite reductase marker genes nirK and nirS were more found in the pristine sediment than the polluted site. The microorganisms containing this marker gene are more diverse and were distributed at both sampling sites. At S1 and S6, 13% and 7% of all nirK reads were mapped to Thaumarchaeota, while no reads of nirS were affiliated to archaea. At the pristine site 23% and 4% of the reads mapped to uncultured bacteria for nirS and nirK respectively, the number of reads mapped to this group decreased in S6. A higher abundance was found for nitric oxide reductase (*norB/norZ*) reference database, with 26 and 18% of all reads matching to *norB* and *nosZ* at S1 respectively, and 23 and 15% at S6. The marker genes for anammox, hydrazine syntheses (hzsA) and hydrazine dehydrogenase (hdh), were not found in the metagenomes. The key gene for dissimilatory nitrate reduction to ammonium (DNRA) is *nrfA* (encoding for the penta-heme nitrite reductase), which had more potential at S6 (16%) compared to S1 (9%).

The nitrogen fixation potential was found with the use of the marker gene *nifH* (encoding for a subunit of the nitrogenase). In total 47 and 58 reads matched *nifH* in S1 and S6 respectively. The gene was more found in Archaea in S6 compared to S1, while the potential bacterial community was diverse.

#### Methane cycling potential

To find the methane cycling potential in the Citarum river sediment marker genes were used targeting methane production and oxidation. The gene encoding for methyl-coenzyme M reductase (mcrA) was used to find the potential for methane production and anaerobic methane oxidation. At S1 12 reads matched to mcrA, which were mostly related to uncultured archaeon, 4 reads were related to ‘*Candidatus* Methanoperedens nitroreducens’, and one hit was affiliated to an uncultured methanogen. While a higher abundance was found at S6 (56 reads), where 14% of the reads were affiliated to *Methanobacteriales* and 86% of the reads to methanogens. For the aerobic methane oxidation, the marker genes *pmoA* and *mmoX* (encoding particulate and soluble methane monooxygenase) were used and 8 reads were found in the dataset of S1 where 2 reads were affiliated to type I methanotrophs while rest of the reads was uncultured.

## DISCUSSION

The effect of river pollutants on the microbial communities in sediments is a challenge to unravel because of the many possible interactions that can occur. The present study provided an insight into the microbial diversity in the sediment of the upper Citarum river basin, which is highly impacted by inputs from agricultural activities, industry and urban run-off. Sediment samples were collected along a gradient of pollution, from the pristine source of the river to the heavily polluted downstream site in the densely populated urban area of Bandung. From the 16S rRNA gene amplicon sequencing, differences in microbial diversity and community structure were observed with increasing pollution. While the Simpson index did not show much variation, the Shannon diversity index and Shannon evenness decreased in number, indicating that the diversity decreased and that the microbial community structure becomes less evenly distributed along the river. These differences in microbial community distribution was also observed with the phylogenetic classification based on the 16S rRNA gene read retrieved from metagenomics datasets. Along the gradient of pollution this evenness even decreases further, but not to high extend, indicating that the pollution at Cisangkuy (S3) already has a great influence on the survival and community distribution of the microbial community. Similar results were found by Saxena *et al.* (2018), where microbial community diversity profiles decreased with increased exposure to urban and industrial pollution.

The major bacterial phylum at all sites, where genomic DNA was extracted and analyzed, were the *Proteobacteria*, with dominance of *Beta*- and *Deltaproteobacteria* at the source of the river (S1 and S2), which shifted to relative high abundance of *Alpha*- and *Gammaproteobacteria* as soon as the river became polluted (S3) till the polluted downstream sites (S6). In several freshwater systems the *Betaproteobacteria* were the most abundant group (Böckelmann *et al.*, 2000; Battin *et al.*, 2001; Liu *et al.*, 2012), while other studies found a decrease in there occurrence with urban pollution (Saxena *et al.*, 2018). Other abundant phyla at the pristine sites were the Acidobacteria, Cyanobacteria and Nitrospirae. Acidobaceria are versatile heterotrophs and are capable of nitrate and nitrite reduction, while most Nitrospirae are versatile aerobes specialized in oxidation of nitrite to nitrate. This corresponds with the slight increase in nitrification marker genes found at S1 compared to S6. Bacterial phyla that were highly abundant at the polluted sites were *Bacteriodetes*, which were also found in studies of the Zenne River in Belgium (García-Armisen *et al.*, 2014), Athabasca River in Canada (Yergeau *et al.*, 2012), and in Jaboatão River in Brazil (Kochling *et al.*, 2017) which are all rich in organic matter. *Bacteriodetes*, are implicated in the degradation of complex organic substrates (Kabisch *et al.*, 2014). The members of *Actinobacteria* are often found in polluted waters (Yergeau *et al.*, 2012; García-Armisen *et al.*, 2014), where they potentially play an important role in the breakdown and recycling of organic compounds. Similarly, *Clostridiales* were abundant at S3 to S6 and are able to degrade very recalcitrant organic compounds. *Clostridiales* are also indicators for feacal contamination, that could come in the water due to live stock manure and suboptimal sewage systems that discharge effluents in the river. In addition, the presence of *Lactobacillales* at S6 also indicates the release of sewage in the river after Bandung city. The relative abundance of the archaea was similar in all sediments, *Soil Crenarchaeotic group* had the highest relative abundance at all sites (between 28-79%). They have been found in many different ecosystems like forest soils, agricultural sites and contaminated soils (Sandaa *et al.*, 1999; Ochsenreiter *et al.*, 2003). There are indications that these soil Crenarchaeota may contribute to ammonia oxidation, but whether they contribute to the nitrogen cycle in these sediments would need further study (Treusch *et al.*, 2005; Ayton *et al.*, 2010). Of the other abundant archaea, the methanogen abundance became more prominent with increasing pollution as also reported by Kimes et al (2013) in sediment samples of the Gulf of Mexico. This was corroborated by the increase of the methanogenic marker gene *mcrA* at the polluted site, indicating that more fermentation products (i.e. hydrogen, or acetate) are available to the methanogens in the polluted sites. The change in diversity between the source and polluted downstream sites of the river are indications that the sediment changed from oxic to anoxic environment.

The genomic nitrogen cycling potential in the river sediment was in general low. In this analysis we corrected for sequencing depth and gene length, however we could not correct for gene copy number since for many microorganisms this is not yet known, for example the gene *hao* can have up to ten different copies in anammox bacteria (Van de Vossenberg *et al.*, 2013; Speth *et al.*, 2017). At the pristine site ammonium oxidation (*amoA, nxr*) and denitrification genes (*nar, nir, nor, nos*) were more abundant N-cycle processes, whereas ammonification (*nrf*) seemed more important in S6. Interestingly, N-fixation potential (*nifH*) could be found in all sites, while anammox (*hzsA, hdh*) could not be detected. The absence of anammox was unexpected since it has been found before in freshwater river sediments (Zhao *et al.*, 2013; Lisa *et al.*, 2014). The methane cycle genes *pmoA* and *mcrA* were much less covered and showed a higher potential for methanogenesis at the polluted site.

In conclusion, we showed that with the increase of pollution in the Citarum river, the microbial community composition decreased in evenness and the relative abundance of *Alpha*- and *Gammaproteobactria, Clostridia*, and *Actinobacteria* increased. The potential for nitrogen turnover is relatively low in this river system. Since the sediment becomes more anoxic at the polluted site, methanogenesis potential is increased compared to the source of the river. Increased wastewater treatment and source separation would help to improve water quality and restore diverse microbial communities.

## ACKNOWLEDGMENTS

We thank the sampling team for the great help during the sample trip in Bandung, Indonesia, and Theo van Alen for all the help with metagenomic sequencing. This research was funded by the Netherlands Organization for Scientific Research through the Netherlands Earth System Science Center (NESSC) Gravitation Grant [grant number 024.002.001 to AdJ, MitZ and MJ], the Soehngen Institute of Anaerobic Microbiology (SIAM) Gravitation Grant [grant number 024.002.002 to MJ] and the European Research Council Advanced Grant [grant number 339880 to MJ].

## SUPPLEMENTAIRY DATA

**Supplementary Figure 1.**
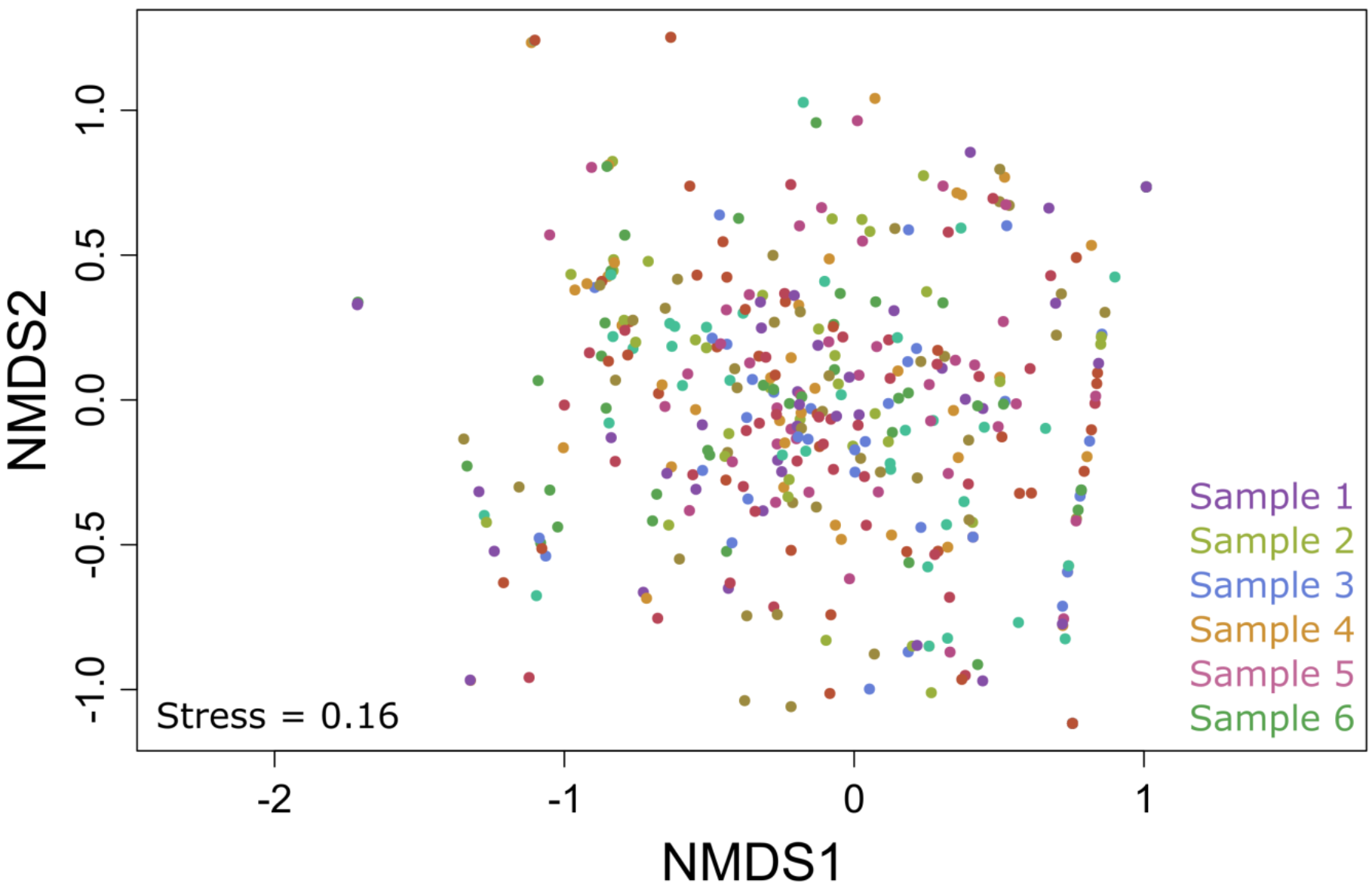
Non-metric multidimensional scaling (NMDS) plot for the archaeal community structure on OTU level for sample 1 to 6, with an average stress level of 0.16 using 20 iterations.

**Supplementary Figure S2.**
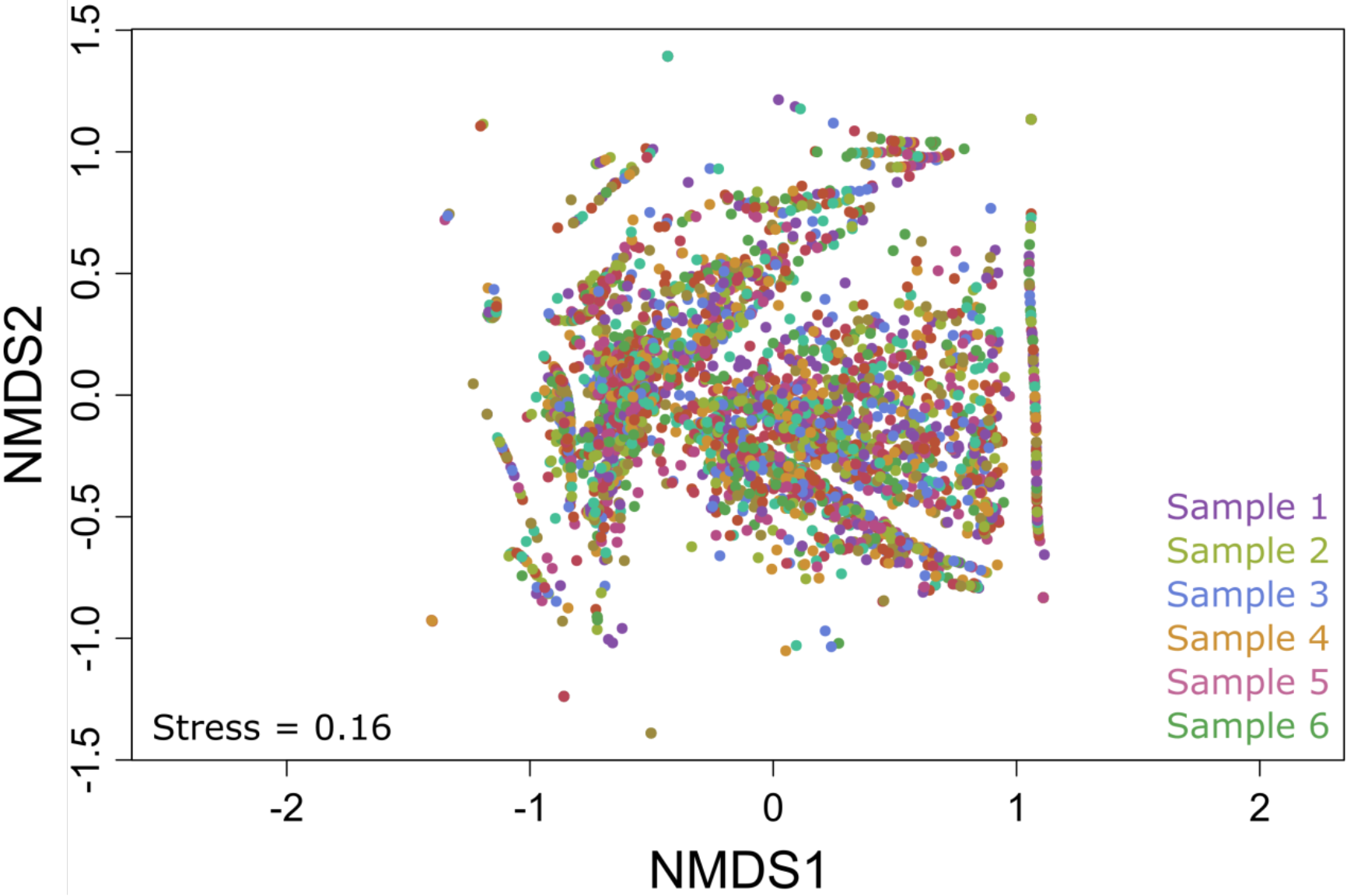
Non-metric multidimensional scaling (NMDS) plot for the bacterial community structure on OTU level for sample 1 to 6, with an average stress level of 0.16 using 20 iterations.

**Figure S3.**
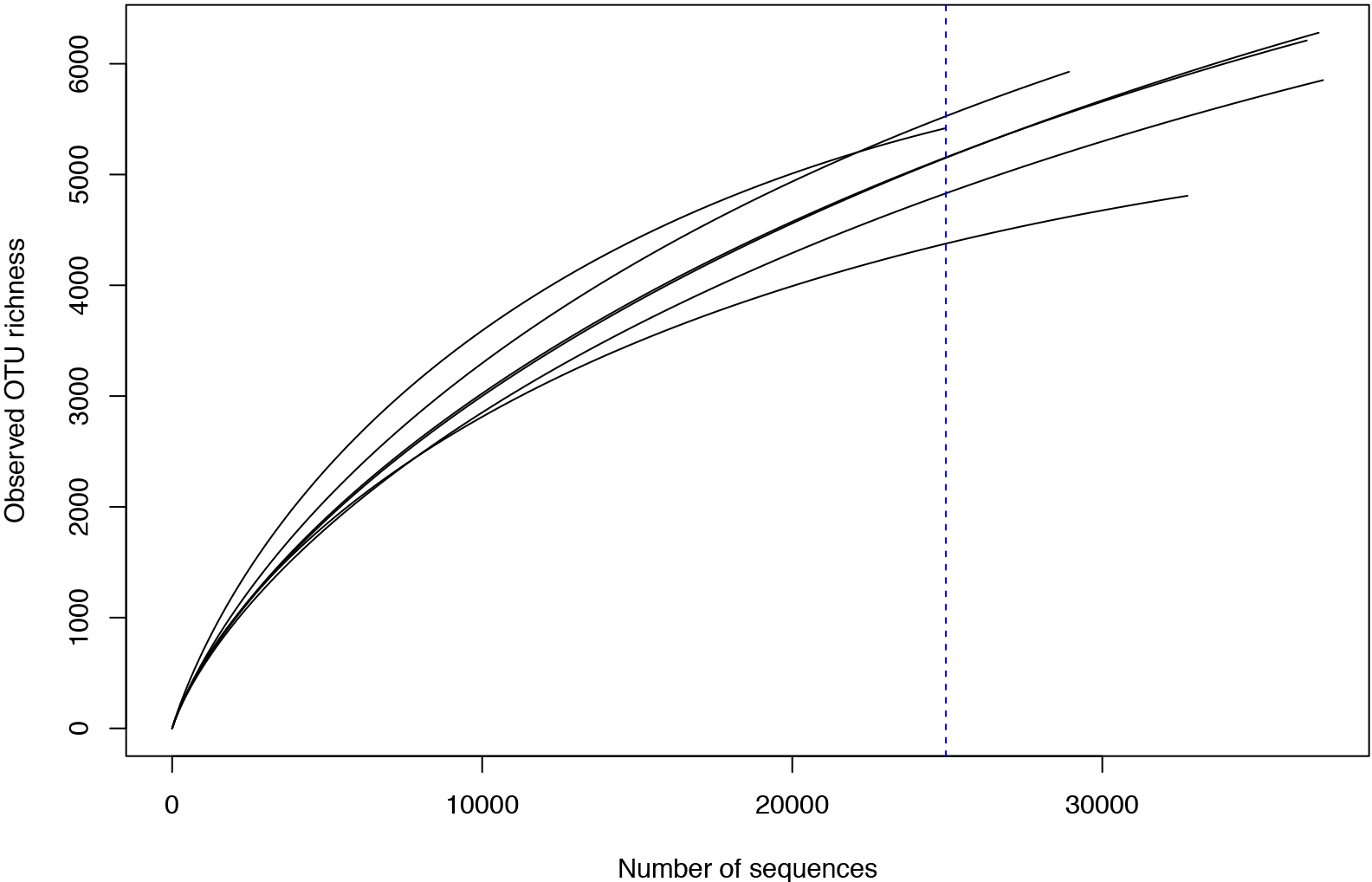
Rarefraction curves of bacterial 16S rRNA gene amplicon sequences after singleton removal.

**Figure S4.**
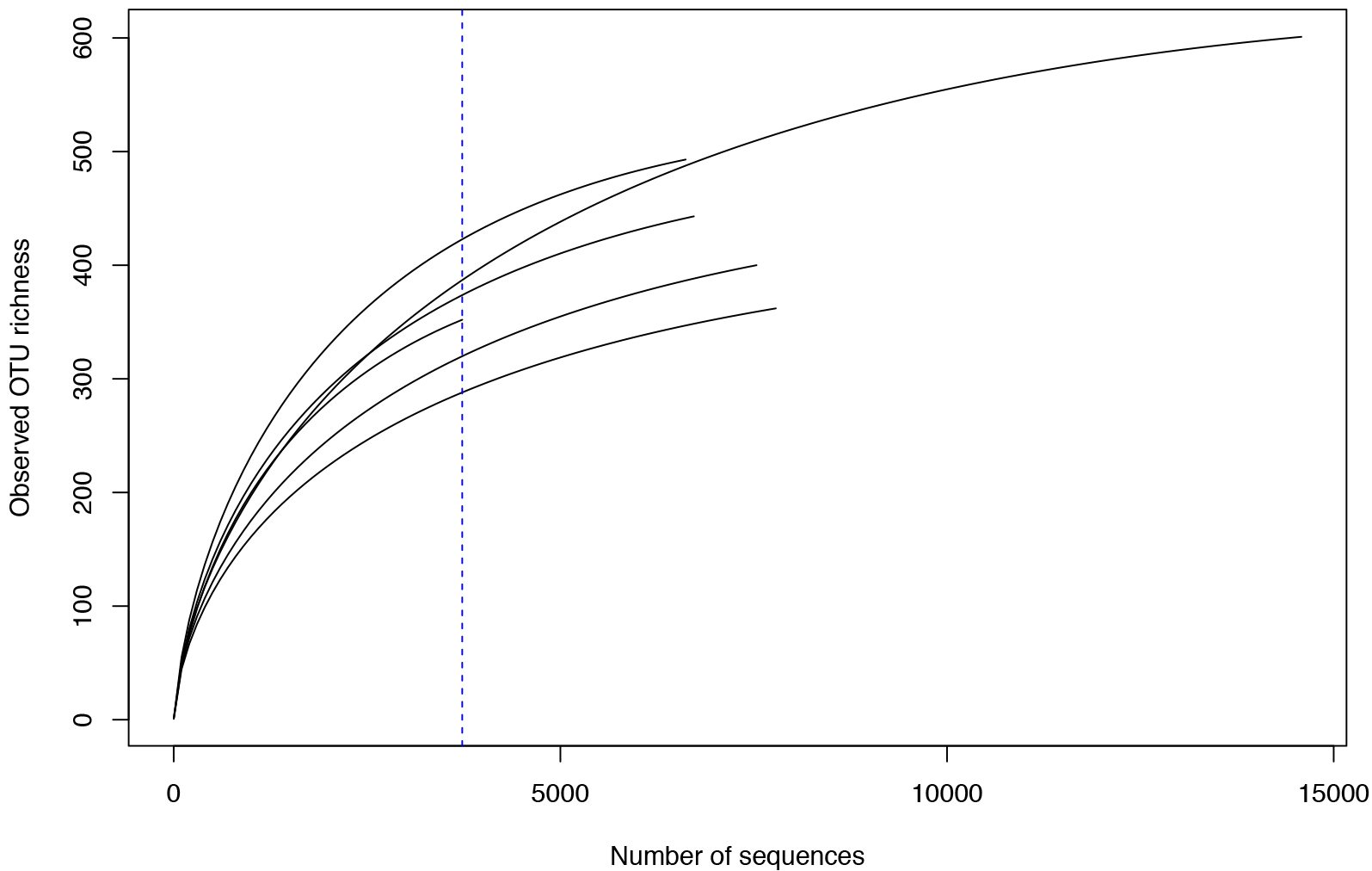
Rarefraction curves of archaeal 16S rRNA gene amplicon sequences after singleton removal.

**Table S1.** Data from the Citarum metagenomic libraries described in this study.

| Features | S1 | S6 |
| --- | --- | --- |
| Individual reads prior to QC | 2.943.692 | 2.129.763 |
| Average length of reads prior to QC (bp) | 246.5 | 255.9 |
| Percentage of reads that were trimmed | 87.76 | 89.90 |
| Individual reads post trimming | 2.583.249 | 1.914.762 |
| Average length of reads post trimming (bp) | 268.3 | 271.9 |
| Mapped reads against SILVA database | 3.076 | 3.017 |
| Individual reads after blast against SILVA database | 2.266 | 2.481 |
| Aligned reads against SILVA database | 2.204 | 2.381 |

